# Nilotinib activates endothelial TLR4 to exacerbate atherosclerosis

**DOI:** 10.1101/2019.12.14.876599

**Authors:** Dan Qu, Mingyu Huo, Li Wang, Chi-Wai Lau, Xiao Yu Tian, Yu Huang

## Abstract

The use of nilotinib (Tasigna®), a second-generation tyrosine kinase inhibitor for treating chronic myeloid leukemia, increases risks for atherosclerosis. Here, we demonstrate that in endothelial cells, nilotinib activated TLR4, triggerd expression of inflammatory molecules, and increased monocyte attachment, which were all inhibited by knockdown of TLR4 or TLR4 inhibitor, CLI-095. Orally administered nilotinib profoundly accelerated atherosclerotic lesion formation in ApoE^−/−^ mice, while co-administration of CLI-095 effectively reduced lesion areas. Our findings reveal TLR4 activation as an underlying mechanism of the pro-atherosclerotic effect of nilotinib and suggest TLR4 inhibition as an effective therapeutic approach to address vascular safety issue of nilotinib.

## Introduction

Cardiovascular complication is one of the critical safety concerns associated with the use of tyrosine kinase inhibitors (TKI) for cancer therapy(1–4). Nilotinib (Tasigna®), a second-generation BCR-ABL TKI, is an alternative to the first-generation BCR-ABL inhibitor imatinib for Philadelphia chromosome positive chronic myelogenous leukemia (Ph+ CML) patients who are resistant or intolerant to imatinib. Nilotinib was approved by FDA as first and only CML therapy with Treatment-free Remission data in its label in 2017. Though nilotinib has achieved good tolerability and high potency, the emerging vascular adverse effects have limited its use as front-line therapy for CML(5–9). Here, we aimed to investigate the cellular mechanisms involved and to identify a therapeutic strategy to alleviate nilotinib-exacerbated atherosclerosis.

Nilotinib works to limit the excessively production of leukocytes. The resulting low level of circulating leukocytes, as we observed in the nilotinib-treat mice (Supplemental Figure S3), makes leukocytes unlikely to contribute to atherogenesis. On the other hand, endothelial cells (ECs) serve as the front barrier of the vasculature and govern the health of blood vessels(10). Endothelial inflammation is the initial pathogenic event of atherosclerosis and an early marker that precedes angiographic or ultrasonic evidence of structural emergence of atherosclerotic lesions(11). Therefore, the present study focuses on the effect of nilotinib on endothelial cells during atherogenesis.

Toll-like receptor 4 (TLR4) not only detects foreign pathogens, but also participates in sterile inflammation including atherosclerosis(12). TLR4 expression is elevated in atherosclerotic lesions, preferentially in ECs and infiltrated macrophages(13–15). Global knockout of TLR4 results in reduced lesions in atherosclerotic models of LDLr^−/−^ and ApoE^−/−^ mice(16–18). Furthermore, we have recently identified TLR4 as a sensor for disturbed flow and mediated disturbed flow-induced endothelial inflammation and accelerated atherogenesis(19). Given the critical role of TLR4 in atherogenesis, we questioned whether TLR4 mediates the vascular adverse effects of nilotinib.

We demonstrated here that nilotinib induced TLR4-dependent endothelial inflammation and greatly exacerbated lesion formation in both acute (partial carotid ligation induced) and chronic atherogenesis model in *ApoE*^*−/−*^ mice. CLI-095, the TLR4 inhibitor, effectively attenuated endothelial inflammation and reversed the nilotinib-exacerbated atherogenesis in *ApoE*^*−/−*^ mice. Our findings suggest TLR4 inhibition as an effective therapeutic approach to address the vascular safety concern of nilotinib.

## Results and Discussion

Nilotinib induced death of human umbilical vein endothelial cells (HUVECs) in a concentration-dependent manner (Fig S1). Nilotinib at 30 μM killed 45% cells in 24 hours, which was prevented by co-treatment with TLR4 inhibitor CLI-095. While nilotinib at 10 μM produced a moderate cytotoxic effect as 75% cells were viable after 24-hour treatment. Ten μM of nilotinib is near the clinically observed peak serum concentration(20–22), therefore this concentration was used for the following experiments.

TLR4 signaling was mediated through MyD88-dependent pathway in HUVECs(19, 23). Proximity ligation assay showed that nilotinib induced interaction of MyD88 to TLR4 (Fig 1A), indicating activation of TLR4 signaling. The expression of downstream inflammatory genes, VCAM1 and CCL2, were also induced by nilotinib (Fig 1B, C), which consequentially led to recruitment of monocytes to HUVECs (Fig 1D). Such inflammatory responses were reversed by TLR4 inhibition with CLI-095 or knockdown of TLR4 (Fig 1B-D).

**Figure 1.**
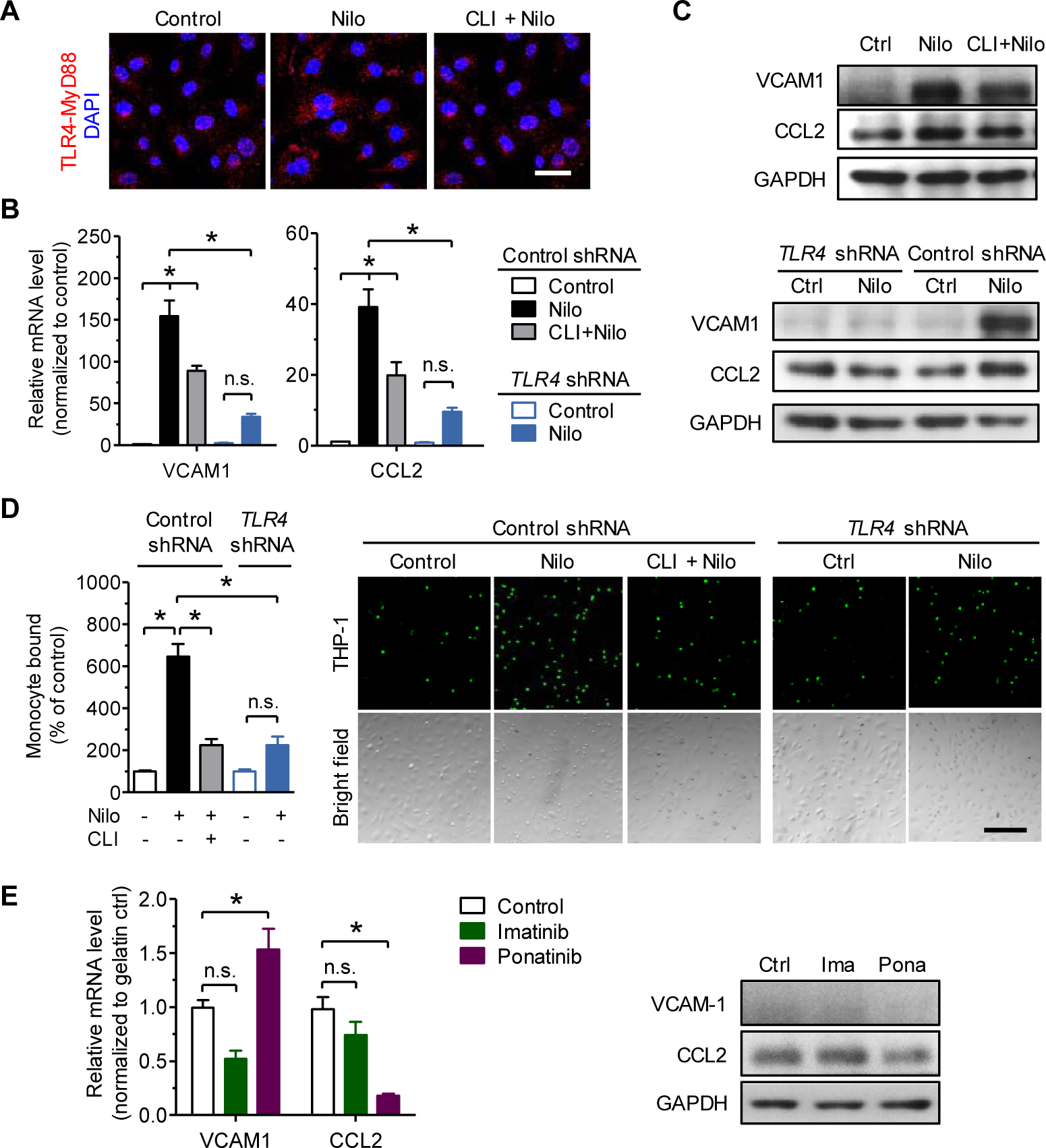
TLR4 mediates nilotinib-induced endothelial inflammation. **A**-**C**, HUVECs were treated with nilotinib (10 μM) and CLI-095 (100 nM) for 4 hours and analyzed for TLR4-MyD88 interaction (**A**, n = 4, scale bar: 50 μm) and the expression of inflammatory makers (**B** and **C**, n = 4). **D**, HUVECs were treated with indicated drugs for 4 hours, washed to remove drugs and assayed for monocytes attachment (THP-1, green fluorescence) (n = 5 for control shRNA; n = 4 for *TLR4* shRNA). Scale bar: 200 μm. **E**, HUVECs were treated with Imatinib (10 μM) and Ponatinib (1 μM) for 4 hours and analyzed for expression of inflammatory makers, n = 5. Data are presented as mean ± SEM. \**P* < 0.05 by one-way ANOVA with Tukey post-test (**B**, **D**, and **E**). Nilo, nilotinib; CLI, CLI-095.

Imatinib (Gleevec) is the first-generation BCR-ABL TKI, with a relatively safe profile(8, 24). imatinib at 10 μM, close to its peak serum concentration, was not toxic to ECs (Fig S2) and hardly induced expression of inflammatory genes VCAM1 and CCL2 (Fig 1E). On the other hand, the third-generation TKI ponatinib (ICLUSIG) has been warned for cardiovascular adverse events(25). Ponatinib at 1 μM (close to its peak serum concentration) killed 90% ECs in 24 hours. Ponatinib induced a slight increase of VCAM1 mRNA expression (Fig 1E), which was negligible as compared to that induced by nilotinib. Therefore, imatinib and ponatinib at clinically relevant concentrations failed to reproduce the pro-inflammatory effect of nilotnib in ECs (Fig 1E), indicating that the pro-inflammatory effect in ECs is specific to nilotinib. The cytotoxic effect of nilotinib may also be different from the direct cytotoxic effect of ponatinib.

Nilotinib-treated patients tend to develop peripheral arterial occlusive diseases, cerebral ischemia, and myocardial infarction(6). To confirm the pro-atherosclerotic effect of nilotinib, we treated ApoE^−/−^ mice with 8 mg/kg nilotinib by oral administration twice daily. This dosage is equivalent to that prescribed to CML patients (400 mg b.i.d.). After two months, nilotinib exacerbated atherosclerotic lesion coverage to ~18% compared to control of ~9% (Fig 2A). Notably, CLI-095 treatment reduced the lesion coverage to ~14%. In addition, nilotinib treatment reduced the number of total circulating white blood cells and monocytes (Fig S3). Therefore, leukocytes are unlikely contribute to nilotinb-exacerbated atherogenesis because they did not increase after nilotinib treatment. This observation is in line with a latest publication that nilotinib induces necrosis in monocytes and inhibits their differentiation into macrophages(26).

**Figure 2.**
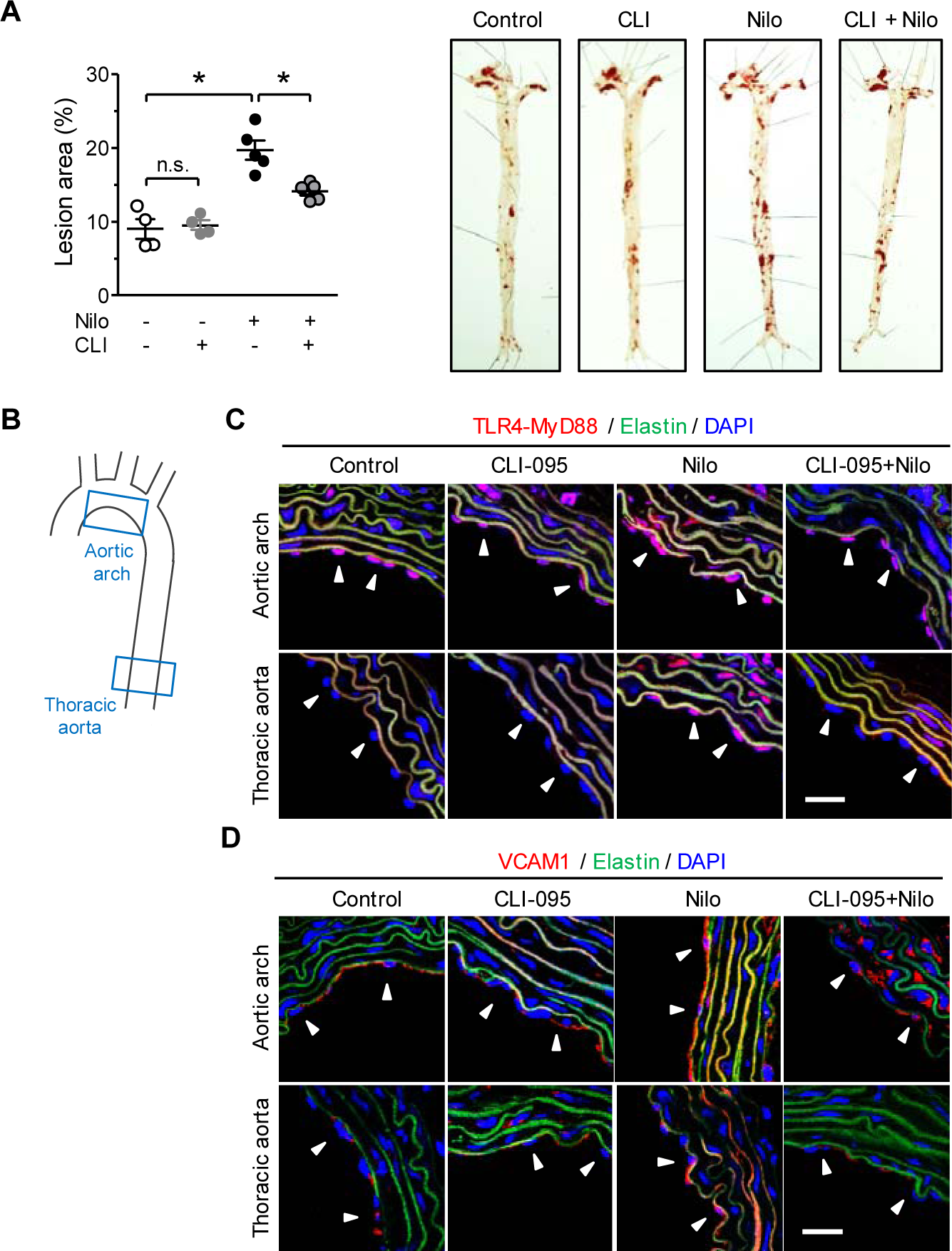
CLI-095 reverses nilotinib-induced atherosclerosis. **A**, ApoE^−/−^ mice treated with nilotinib (8 mg/kg, twice per day, p.o.) and/or CLI-095 (0.1 mg/kg/day, i.p.) for two months. Lesion areas were stained by oil-red and analyzed (n = 4-5 mice for each group). **B**, The schematic diagram shows areas of mouse aorta examined by immunofluorescence staining. **C** and **D**, C57BL/6J mice were treated with nilotinib and/or CLI-095 (same dosages as for ApoE^−/−^ mice) for three days. Cross section of aortic arch and thoracic aorta were stained for TLR4-MyD88 interaction (**C**) and VCAM1(**D**). Arrowheads indicate endothelial cells on the luminal side of vascular wall. Green was autofluorescence of elastin. Nuclei were counterstained with DAPI. Images are representative of four mice. Scale bar: 20 µm. Nilo, nilotinib; CLI, CLI-095.

We next examined the TLR4 activation *in vivo* in the endothelium of mouse aortas (Fig 2B-D). As the fatty streak and plque formation in *ApoE*^*−/−*^ mice interfered with the examination of endothelium, C57BL/6J mice were used instead and treated with nilotinib and/or CLI-095. Atherosclerotic lesions develop preferentially at arterial branches and curvatures in contrast to relatively straight thoracic aorta(27). Therefore, the curved aortic arch was carefully distinguished from thoracic aorta. The basal TLR4 activation of MyD88 and VCAM1 expression levels were higher in aortic arch compared to those in thoracic aorta, as shown in the control group (Fig 2C, D). While nilotinib activated TLR4 signaling in thoracic aorta and further augmented it in aortic arch. CLI-095 treatment reversed TLR4 signaling to its basal level in both aortic arch and thoracic aorta. This result is consistent with that CLI-095 effectively inhibited nilotinib-exacerbated atherogenesis in *ApoE*^*−/−*^ (Fig 2A).

To further confirm the above findings, we employed the partial carotid ligation model of accelarated lesion formation (Fig 3A)(28, 29). Left carotid artery (LCA) of *ApoE*^*−/−*^ mice was partially ligated and within two weeks atherosclerotic lesions developed to 21% of total areas (Fig 3B, C). The lesion area was exacerbated to 48% by nilotinib treatment, which was reduced to 13% by CLI-095. In contrast, nilotinib did not induce any lesion formation in the intact right carotid arteries (RCAs) during the treatment.

**Figure 3.**
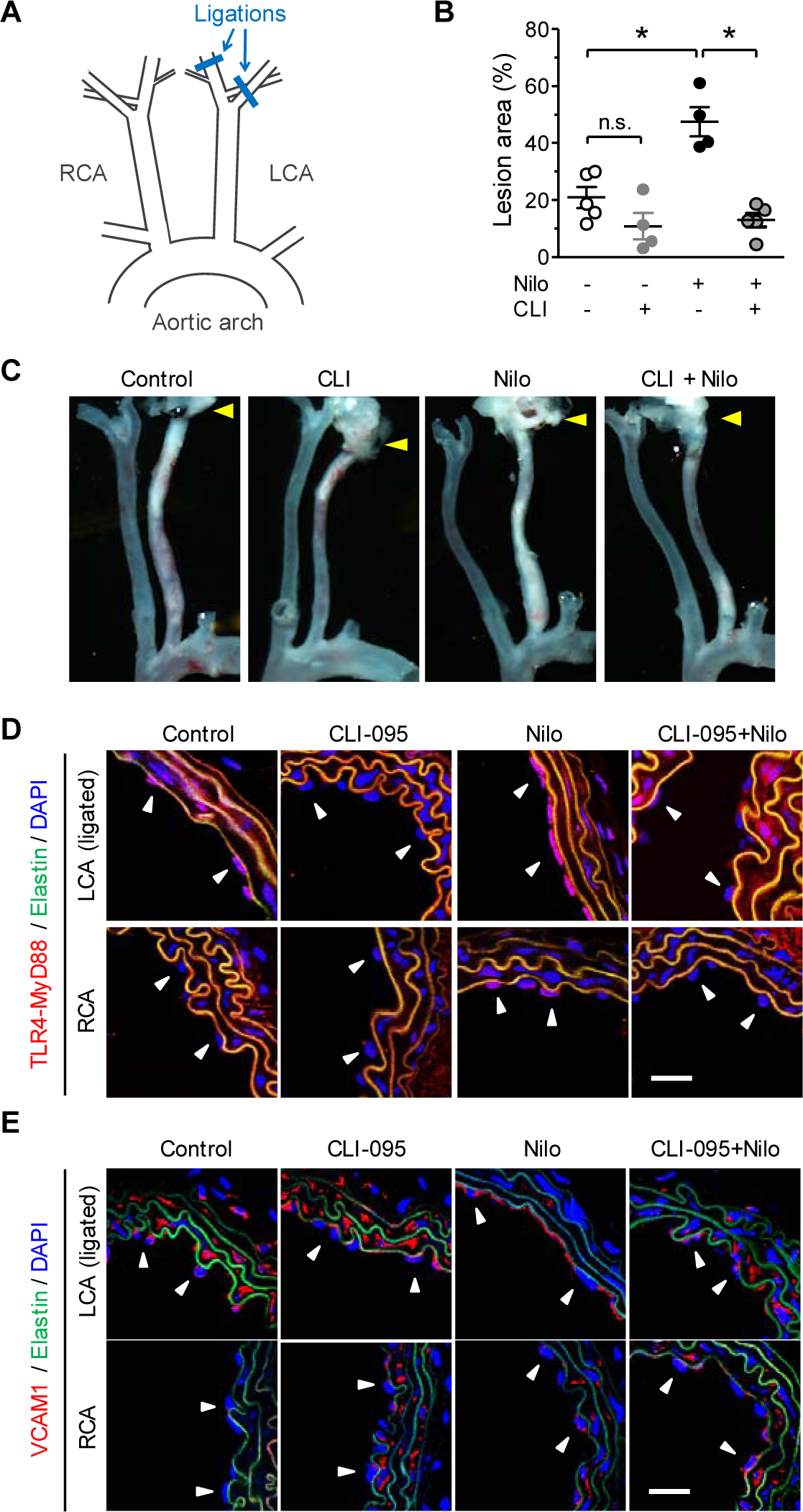
CLI-095 reverses nilotinib-induced atherosclerosis and TLR4 activation. **A**, The schematic diagram depicts partial ligation of left carotid artery (LCA) that acutely induces atherosclerosis. The right carotid artery (RCA) is left intact. **B** and **C**, ApoE^−/−^ mice underwent partial ligation were treated with nilotinib (8 mg/kg, twice per day, p.o.) and/or CLI-095 (0.2 mg/kg/day, i.p.) for two weeks. Lesion areas in LCA were analyzed. Arrowhead indicates the ligations on branches of LCA, n = 4-5. **D** and **E**, C57BL/6J mice underwent partial carotid ligation and were treated with respective drugs for three days. Cross section of carotid were stained for TLR4-MyD88 interaction and VCAM1 expression. Arrowheads indicate endothelial cells on the luminal side of vascular wall. Green was autofluorescence of elastin. Nuclei were counterstained with DAPI. Images are representative of four mice. Scale bar: 20 μm. Nilo, nilotinib.

In order to examine the *in vivo* effect of nilotinib and CLI-095 on ECs, the partial carotid ligation experiment was replicated in C57BL/6J mice. Similar to our observation in chronic atherosclerosis model, nilotinib augmented TLR4 activation and VCAM1 expression in LCA, which were reduced by CLI-095 (Fig 3D, E). Notably, in both thoracic aorta and RCA, nilotinib induced TLR4 activation, albeit to a lesser extent. Yet this minor TLR4 activation in RCA did not lead to a significant production of VCAM1 nor accelerated lesion formation within the treatment period (Fig 3D,E). This is probably due to the protective effect of laminar flow to suppress TLR4-dependent inflammation(19) and hence diminishes the adverse effect of nilotinib within the course of the treatment. Taken together, the present results strongly indicate that targeting TLR4 could be a promising approach to mitigate the vascular adverse events of nilotinib.

Though nilotinib was designed to specifically inhibit tyrosine kinase activity of BCR-ABL, it off-targets other kinases such as PDGFRα and KIT(30). Nilotinib probably docks in the pocket of MD2 and interacts with TLR4, mimiking the LPS/TLR4/MD2 complex upon TLR4 activation by LPS. Further investigation is required to understand the molecular mechanism. Although it was unclear whether nilotinib acted on endothelial cells through off-target activation of TLR4 or other non-specific effects, the pro-atheroslcerotic effect was reversed when TLR4 was knocked down or inhibited by CLI-095. These results suggest, for the first time, a potential therapeutic value of TLR4 inhibitors for CML patients to whom nilotinib is their best choice.

Several TLR4 antagonists have been evaluated in clinical trials for treatment of sepsis and inflammatory diseases (31). For instance, eritoran and TAK-242 (CLI-095) underwent Phase III clinical trial for sepsis; yet the former did not reduce mortality and the latter was suspended during the trial (31, 32). Fortunately, none of studies or clinical trials on these inhibitors have reported drug safety concerns. Hopefully, re-purpose of TLR4 inhibitors for atherosclerosis will help to ameliorate vascular adverse events of nilotinib.

## Methods

Detailed experimental methods are included with the supplemental materials.

### Study approval

C57BL/6J and ApoE^−/−^ mice were supplied by the Univeristy Laboratory Animal Services Center, the Chinses University of Hong Kong. The use of animals was approved by the Animal Research Ethical Committee, Chinese University of Hong Kong (17-050-MIS) and conformed to the Guide for the Care and Use of Laboratory Animals published by the US National Institute of Health (NIH Publication No.85-23, revised 1996).

## Acknowledgments

This study was supported by National Science Foundation of China (91339117), Research Grants Council of Hong Kong (C4024-16W, CUHK14105814, CUHK464712, C7055-14G), Hong Kong Croucher Foundation, and CUHK Vice-Chancellor’s Discretionary Fund.

## Author contribution

DQ designed the project, performed most experiments, analyzed data, and wrote the manuscript. MH and CWL assisted with experiments. LW and XYT adviced on the project design. LW, XYT, and YH revised the manuscript. YH provided funding and supervised this project.

