## supplemental materials for "Nilotinib activates endothelial TLR4 to exacerbate atherosclerosis"

### Materials and Methods

*Cell culture*

Human umbilical vein endothelial cells (HUVECs) were cultured in 0.1% gelatin-coated flasks with EGM-2 (Lonza, Walkerville, Maryland, USA). Passage 5-8 cells were used in all experiments. THP-1 cells were cultured with DMEM supplemented with 10% FBS and 1% penicillin/streptomycin. All cell lines were purchased from American Type Culture Collection (UA, USA) and culture materials were purchased from Gibco (Thermo Fisher Scientific, Waltham, WA, USA). The cell viability was assessed by WST-1 assay (Sigma-Aldrich, St. Louis, MO, USA) according to the manufacturer`s instruction using a microplate reader (xMark microplate spectrophotometet, Bio-Rad, Hercules, CA, USA). For monocyte adhesion assay, HUVECs were treated and washed to remove drugs, then incubated with THP-1 ($5x10^5$ cells/ml, fluorescence-labelled with Calcein-AM, Thermo Fisher Scientific) for 30 minutes at 37°C. Non-adherent monocytes were gently washed with DMEM. The number of adherent monocytes was counted in 200x magnification fields.

*Animal studies*

All mice used were male and aged at 12-14 weeks at the beginning of treatment. Mice were housed in animal holding rooms at room temperature and under 12-hour light/dark cycle, and fed standard chow and water *ad libitum*, unless specified. To obtain mouse arteries for *ex vivo* studies, mice were euthanized by $CO_2$ inhalation. Aortae or carotid arteries were excised, cleaned of adhering tissues, and prepared for immunofluorescence study.

*Chronic atherosclerosis model*

12-week-old $ApoE^{-/-}$ (fed on western diet (0.21% cholesterol, Research Diet, New Brunswick, NJ, USA)) or C57BL/6J mice (fed on normal chow) were grouped to receive both/either/neither nilotinib (Sigma-Aldrich) and/or/nor CLI-095 (Invivogen, CA, USA). 8 mg/kg Nilotinib (or equivoluminal water for non-nilotinib groups) was given twice per day by oral administration. Mice were starved 2 hours before and 1 hour after the drug administration according to the instruction to human patients. 0.2 mg/kg/day CLI-095 (or equivoluminal PBS for non-CLI-095 groups) was injected intraperitoneally before the starvation in the morning. After 3-day treatment, C57BL/6J mice were sacrificed and the aortae were dissected out for immunofluorescence staining. After 10-week treatment, $ApoE^{-/-}$ mice were sacrificed and the aortae were dissected out and atherosclerotic lesion area was measured by *en face* Oil Red O staining of whole aorta. Total surface area and lesion areas in the aortae were quantified by ImageJ software (National Institute of Health, Bethesda, MD, USA).

*Partial carotid ligation*

Partial carotid ligation (1) was employed to acutely induce atherogenesis in C57BL/6J and $ApoE^{-/-}$ mice aged at 12 weeks. Mice were anesthetized by intraperitoneal injection of xylazine (10 mg/kg) and ketamine (80 mg/kg) mixture. The ventral cervical skin was depilated and disinfected with 75% ethanol. A vertical incision (5–7mm) was made along the midline of the neck, and the left carotid artery (LCA) was exposed by blunt dissection. The left external carotid, internal carotid, and occipital artery were ligated with 6–0 silk suture (Pearsalls limited, UK), while the superior thyroid artery was left intact. The incision was then approximated and closed. Mice were monitored until recovery in a warm chamber.

After the surgery, mice were grouped and treated with nilotinib and/or CLI-095 as described above for chronic atherosclerosis model. After 3-day treatment, C57BL/6J mice were sacrificed and the carotid arteries were dissected out for immunofluorescence staining. After 2-week treatment, $ApoE^{-/-}$ mice were sacrificed and the carotid arteries were dissected out and imaged with a black background using a stereomicroscope connected to a camera (Leica Microsystems, Buffalo Grove, IL, USA). Atherosclerotic areas were analyzed by ImageJ.

*Immunofluorescent staining*

The aortae or carotid arteries was dissected and embedded in Tissue-Tek optimum cutting temperature medium, frozen in liquid nitrogen, cross-sectioned at 10 μm on a cryotome, and stored at −80°C until used. All cells or arteries were fixed in 4% paraformaldehyde containing 0.1% Triton X-100 (Sigma-

Aldrich) for 15 minutes, blocked with 5% donkey serum for 1 hour at room temperature, and incubated with primary antibodies (VCAM1, ab134047, Abcam, Cambridge, UK) overnight at 4°C, followed by the fluorescent secondary antibodies (AlexaFluor, Thermo Fisher Scientific). Nuclei were counterstained with DAPI. The target proteins were visualized by Fluoview FV1000 laser scanning confocal system (Olympus, Tokyo, Japan).

For proximity ligation assay, Duolink *in situ* reagents (Sigma-Aldrich) was employed and staining was performed according to the manufacturer's instruction. Cells and arteries were fixed as described above and blocked with Duolink Blocking Solution for 30 minutes at room temperature. Primary antibodies (TLR4, AF1478, R&D SYSTEMS, USA; MyD88, ab2064, Abcam) were diluted 1:100 in Duolink Antibody Diluent and used to probe cells overnight at 4°C. After wash, plus and minus PLA probes were added at 1:5 dilution for 2 hours at room temperature, followed by ligation and amplification at 37°C for 30 minutes and 100 minutes, respectively. Nuclei were counterstained with DAPI. Negative controls were performed to exclude background noise or non-specific amplification of the signal.

*Flow cytometry*

Blood samples were taken from ApoE$_{-/-}$ mice that underwent partial carotid ligation and treated with nilotinib and/or CLI-095; white blood cells were analyzed by flow cytometry as previously described (2). Red blood cells were lysed with RBC lysis buffer (Biolegend, San Diego, CA, USA). Then the cells were pelleted and resuspended in FACS buffer (PBS, 2 mM EDTA, and 5% fetal bovine serum), an aliquot of which was used for immunostaining. Live/Dead Aqua (Thermo Fisher Scientific) was used to determine cell viability. Antibodies used after Fc-block (101320) included CD45 (30-F11; 103132), F4/80 (BM8; 123121, 123110), Ly6c (HK1.4; 128016, 128012), Ly6g (1A8; 35-1276), CD11c (N418; 117327), CD115 (AFS98; 135510, 135506), CD206 (C068C2; 141707), CD8 (53-6.7; 100726) from Biolegend; CD11b (M1/70; 60-0112), CD3 (145-2C11; 50-0031), and CD4 (GK1.5; 65-0042) from Tonbo Biosciences (San Diego, CA, USA); and CD301 (ER-MP23; MCA2392A647) from Bio-Rad. Samples were fixed in FluoroFix buffer (BioLegend) and stored at 4°C until analysis. For intracellular staining, cells were fixed and permeabilized using the Fixation/Permeabilization kit (BD Biosciences, San Jose, CA, USA) per the manufacturer's instructions.

*Quantitative RT-PCR*

Total RNA from HUVECs were extracted by using RNeasy Mini kit (Qiagen, Germany) and converted to cDNA using high-capacity cDNA reverse transcription kit (Applied Biosystems, CA, USA). The real-time PCR analysis was performed on an Applied Biosystems ViiA7, using SYBR select mater mix (Applied Biosystems) and following the standard protocol. The PCR primers used were: *hVCAM1-f,* 5`-CCGGATTGCTGCTCAGATTGGA-3`; *hVCAM1-f,* 5`-AGCGTGGAATTGGTCCCCTCA-3`; *hCCL2-f*, 5`-CAGCCAGATGCAATCAATGCC-3`; *hCCL2-r*, 5`-TGGAATCCTGAACCCACTTCT-3`; *hGAPDH-f,* 5`-GGAGCGAGATCCCTCCAAAAT-3`; *hGAPDH-r*, 5`-GGCTGTTGTCATACTTCTCATGG-3`. The gene expression levels were normalized with the housekeeping gene *GAPDH*.

*Western blotting*

Cells were lysed using RIPA lysis buffer. 10 µg protein in each sample was boiled in loading buffer with 5% β-mercaptoethanol for 10 minutes at 95˚C and loaded for sodium dodecyl sulfate polyacrylamide gel electrophoresis (SDS-PAGE). The resolved protein samples were transferred onto a PVDF membrane (Millipore, Billerica, MA, USA) and blocked with 1% bovine serum albumin at room temperature for 30 minutes. Then the membrane was probed with primary antibodies overnight at 4˚C, followed by the corresponding secondary antibodies at room temperature for 2 hours. Protein expression was detected by ECL reagents (Cell Signaling Technology, MA, USA). All protein expressions were normalized to GAPDH.

*Statistics*

Analysis between three or more groups was done by one-way ANOVA followed by Tukey post-test or two-way ANOVA followed by Bonferroni post-test. All statistical analysis and calculations were performed using Prism version 5 (GraphPad Software, San Diego, CA, USA). P values less than 0.05

were considered statistically significant. Data represented means ± SEM. The number of repeated experiments was indicated in individual figures.

### Supplementary Figures

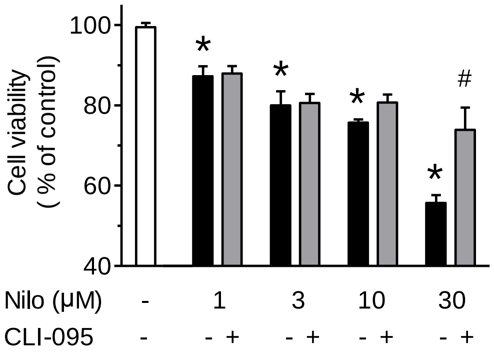


**Figure S1. Nilotinib is toxic to endothelial cells**. HUVECs were treat with 1-30 μM nilotinib, with or without CLI-095 (100 nM), for 24 hours and the cell viability was assessed by WST-1 assay. N = 9. Data are presented as mean ± SEM. **P* < 0.05 vs control, #*P* < 0.05 vs 30 μM Nilo, by Two-way ANOVA with Bonferroni post-test.

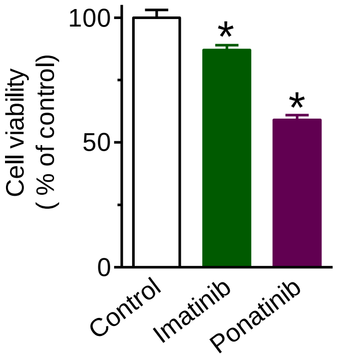


**Figure S2. Cytotoxicity of imatinib and ponatinib.** HUVECs were treat with Imatinib (10 μM) and Ponatinib (1 μM) for 24 hours and the cell viability was assessed by WST-1 assay. N = 8. Data are presented as mean ± SEM. **P* < 0.05 vs control, by Owo-way ANOVA with Dunnett post-test.

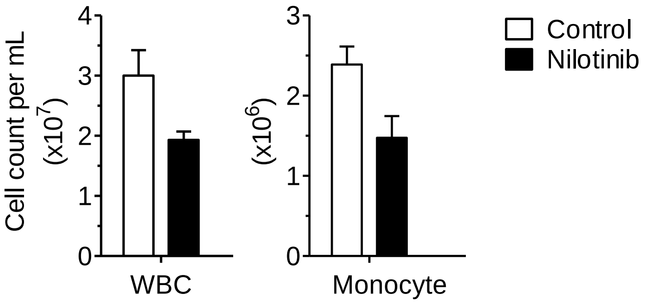


**Figure S3. Nilotinib decreases circulating white blood cells**. $ApoE^{-/-}$ mice were treated with nilotinib and blood were colloected after two weeks. White blood cells were analyzed by flow cytometry. N = 3-4.
